# Splitpea: quantifying protein interaction network rewiring changes due to alternative splicing in cancer

**DOI:** 10.1101/2023.09.04.556262

**Authors:** Ruth Dannenfelser, Vicky Yao

## Abstract

Protein-protein interactions play an essential role in nearly all biological processes, and it has become increasingly clear that in order to better understand the fundamental processes that underlie disease, we must develop a strong understanding of both their context specificity (e.g., tissue-specificity) as well as their dynamic nature (e.g., how they respond to environmental changes). While network-based approaches have found much initial success in the application of protein-protein interactions (PPIs) towards systems-level explorations of biology, they often overlook the fact that large numbers of proteins undergo alternative splicing. Alternative splicing has not only been shown to diversify protein function through the generation of multiple protein isoforms, but also remodel PPIs and affect a wide range diseases, including cancer. Isoform-specific interactions are not well characterized, so we develop a computational approach that uses domain-domain interactions in concert with differential exon usage data from The Cancer Genome Atlas (TCGA) and the Genotype-Tissue Expression project (GTEx). Using this approach, we can characterize PPIs likely disrupted or possibly even increased due to splicing events for individual TCGA cancer patient samples relative to a matched GTEx normal tissue background.

## 1. Introduction

Alternative splicing is a crucial mechanism that underlies the increased complexity of higher eukaryotes. It is now estimated that ∼95% of human genes^1,2^ undergo splicing changes, and the increase in protein diversity that results from splicing has been put forth as one of the primary explanations for the apparent mismatch between species complexity and their genome size.^3,4^ Importantly, alternative isoforms of the same gene can exhibit highly different interaction profiles and thus affect the dynamics of protein interaction networks.^5^ Splicing has been shown to be a key regulator of tissue specificity (especially in the brain),^2,6^ and dysregulation has been increasingly implicated in a wide array of diseases,^7^ from cancer^8,9^ to neurodegenerative diseases.^10^ Thus, it is critical to understand the changes in protein interactions due to splicing that underlie cellular function and dysfunction.

However, a systematic study of splicing-related protein network dynamics is hampered by multiple challenges. Although emergent experimental approaches to directly screen for isoform-level protein-protein interactions are promising,^5^ they are very early in development and highly restricted in resolution. Furthermore, all such screens are naturally bounded by not only a combination of technical and cost constraints, but also the inherent complexity of the underlying networks and the vast number of potential cell types and conditions of interest. Fortunately, the now standard use of RNA-sequencing provides a window into the exploration of splicing patterns across varied conditions. While RNA-seq data alone is still insufficient to chart out the entirety of any particular splicing interaction network, it can be used to understand condition-specific splicing dynamics.

Here, we present Splitpea (SPLicing InTeractions PErsonAlized), a method for detecting sample-specific PPI network rewiring events. Splitpea takes advantage of the key insight that splicing can disrupt critical *protein domains* that mediate PPIs through domain-domain interactions (DDIs), which have been derived based on a mix of structural, evolutionary, and computational approaches.^11–14^ Splitpea integrates PPI and DDI information with sample-specific differential splicing events, and can be used easily in concert with existing, established computational approaches for the identification and quantification of differential splicing.^15^ In the scenario where only an individual sample is available or a different background context is preferable (versus existing control samples), Splitpea provides functionality to use a separate reference database of background splice events; for example, one can choose to use normal GTEx data as background for individual TCGA cancer samples (matched by tissue type). Furthermore, as part of Splitpea’s characterization of the potential downstream interaction network changes, Splitpea indicates likely direction: gain, loss, or chaos (mixed / unclear).

Thus, to our knowledge, Splitpea is the first general tool to characterize potential direction of protein interaction rewiring due to splicing for individual samples. We demonstrate the utility of Splitpea on breast and pancreatic cancer samples from TCGA, using matched normal tissue samples (breast and pancreas) from GTEx. All source code for Splitpea and the corresponding analyses are available via Github (https://github.com/ylaboratory/splitpea), with additional links to download all data and associated networks.

### 1.1 Prior work

Prior work considering domain-domain interactions in the context of splicing have mostly focused on query-based or visualization interfaces. Many consider interactions at the isoform level, aiming to provide a context-specific isoform interaction graph.^16–18^ There has been relatively less work focusing on characterizing network rewiring events. Recently, the first tool to characterize the mechanistic effects of splicing on downstream PPIs was proposed,^19^ but this tool is unable to differentiate between the potential directionality of interaction rewiring (likely gain or loss events). Specifically for the study of cancer, there has also been large-scale analysis efforts to characterize the impact of splicing on PPIs across patients.^8^ Though this work was not patient-specific, it provided strong evidence to demonstrate that there exists a large catalog of isoform changes (with potential downstream impacts on PPIs and regulatory networks) that exist independently of expression changes in cancer. Beyond using PPI networks, there have also been exciting efforts integrating cancer RNA-seq together with somatic mutation data and using functional networks to interpret the downstream impact of splicing.^20^

## 2. Methods

### 2.1 Protein interaction and domain interaction data

Human protein-protein interactions were downloaded from BioGRID (v4.4.207),^21^ DIP (2017-02-05),^22^ HIPPIE (v2.2),^23^ HPRD (Release 9),^24^ Human Interactome (HI-II),^25^ IntAct (2022-04-18),^26^ iRefIndex (v18.0),^27^ and MIPS (Nov 2014).^28^ All proteins were mapped to Entrez Gene IDs.^29^

Known and predicted domain-domain interactions were downloaded from 3did (v2017 06),^11^ DOMINE (v2.0),^12^ IDDI (2011.05.16),^13^ and iPFAM (v1.0).^14^ For predicted DDIs, only interactions with confidence *>* 0.5 were used in downstream analyses.

Protein domain locations were translated to genomic locations using the Ensembl BioMart API and the biomaRt R package^30^ and indexed using tabix^31^ to facilitate fast retrieval given a set of genomic coordinates.

### 2.2 Tissue and tumor splicing data processing

Spliced exon values in the form of percent spliced in (PSI or *ψ*) were obtained for both normal pancreas and breast tissue samples from the Genotype-Tissue Expression (GTEx) project and pancreatic cancer and breast cancer samples from The Cancer Genome Atlas (TCGA) using the IRIS database.^32^ IRIS uses rMATS^33^ to tabulate *ψ* values for skipped exon events (the most abundant splicing event). Though we use rMATS *ψ* values in this study, Splitpea is agnostic to the choice of upstream differential splicing analysis tool and can easily be applied in concert with other tools that use a form of *ψ* as their quantification metric.^34–37^

Specifically, we delineate *ψ*_*i*_ as the *ψ* value for exon *i* = 1, …, *n*_*E*_, where there are *n*_*E*_ total exons that had a reported exon skipping event. Note that the precise exons captured in the sample of interest and the background samples are typically non-identical. We are only able to estimate *ψ* for exons that are captured in both, and thus, *n*_*E*_ represents the number of exons that lie at the intersection of the two larger sets of exons. In the scenario where a background reference distribution of *ψ* values are provided, we calculate ∆*ψ*_*i*_ as the following:

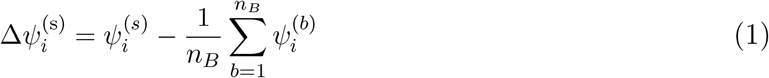

where 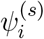is the *ψ* for exon *i* in our sample of interest *s* (e.g., a cancerous pancreatic sample from TCGA), while 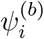 is the *ψ* for the same exon *i* in an individual background sample *b* (e.g., a normal pancreatic sample from GTEx), and *n*_*B*_ is total number of background samples. Intuitively, larger *n*_*B*_ will provide better estimates of the background distribution, especially if there is large variability in splicing patterns. We recommend assembling backgrounds with at least *n*_*B*_ ≥ 30 for the empirical cumulative density function estimate below.

*ψ* values lie in the range [0, 1]; thus ∆*ψ* ∈ [−1, 1], and we are naturally primarily interested in significant events for large |∆*ψ*| values (cases where exons are significantly skipped or significantly retained relative to reference). To calculate an estimated significance level for 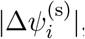, we rely on similar intuition as used in previous studies,^8,38^ that the normal reference samples can be used to construct an empirical cumulative density function for each exon:

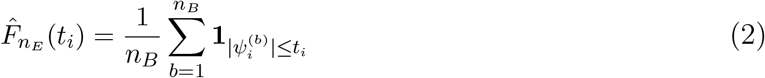

where **1**_**A**_ is the indicator function for event *A*. Given this exon-specific 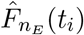, we can estimate an empirical p-value for each exon *i* in sample *s*

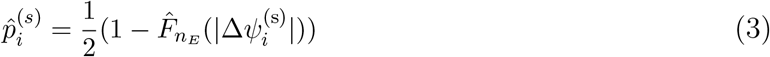

Finally, as input to Splitpea, we filtered exons to only those that are significantly different from background 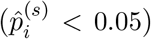 and those with a ∆*ψ* change bigger than 0.05 (|∆*ψ*| *>* 0.05), defined as *ψ* below. We chose to use a p-value cutoff here as opposed to a multiple hypothesis corrected value to reduce false negatives, because we are interested in any possible rewiring events. We hope that this will better enable Splitpea’s use for hypothesis generation tasks. In general, these thresholds can be easily varied depending on the downstream purpose.

### 2.3 Clustering ∆ψ values

For each cancer type, we remove any exons that had missing values in any of the samples, then filtered the exons by variance, keeping only those with variance greater than 0.01. The final set of ∆*ψ* for each cancer type were clustered using the complete hiearchical clustering algorithm and plotted with the heatmap.2 function in the gplots R package.^39^ Clinical annotations for TCGA samples were obtained from the Genomic Data Commons portal with Pam50 calls from Netanely et al.^40^

### 2.4 Network rewiring algorithm

There is inherent complexity in considering the impact of exon changes on protein domains, and finally, proteins, as there are several many-to-many relationships. A single exon can include multiple protein domains, but a single protein domain can also span multiple exons; proteins can thus consist of multiple exons as well as multiple protein domains. Splitpea hones in on potentially domain-mediated protein interactions by first overlaying DDIs on the aggregated PPI network based on the presence of each of the domains that constitute the pair of interactors in the protein. In other words, for a pair of proteins *g*_1_ and *g*_2_, we consider protein domain *d*_1_ in *g*_1_ and domain *d*_2_ in *g*_2_ as potentially mediating a known PPI between *g*_1_ and *g*_2_ if a DDI has been reported between *d*_1_ and *d*_2_. Fig. 1A depicts an example interaction where several DDIs potentially mediate the same PPI.

**Fig. 1.**
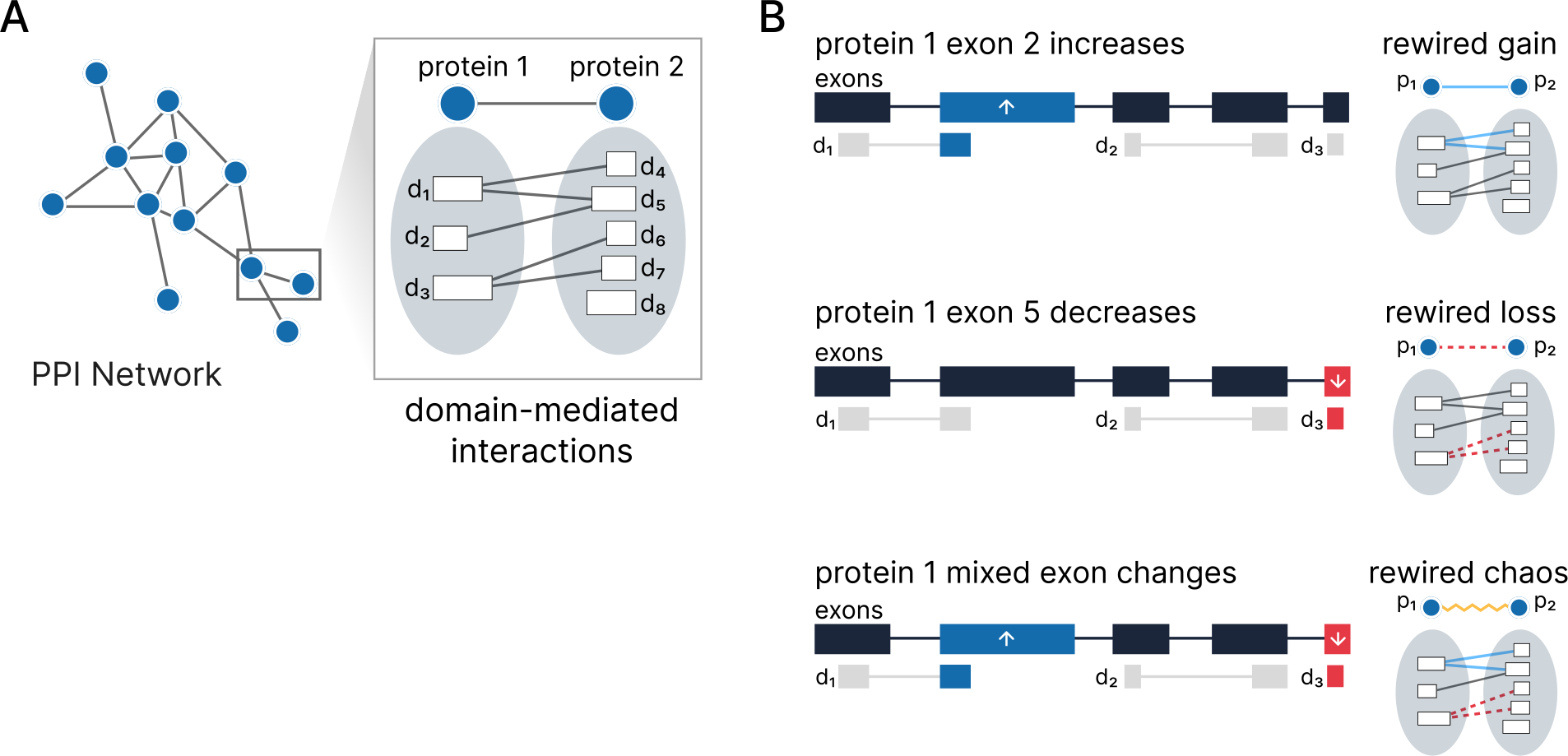
Overview of Splitpea. Splitpea combines prior knowledge in the form of protein-protein and domain-domain interactions with splicing changes to provide a view of a rewired network for a given experimental context. Splitpea defines a rewiring event when exon changes affect an underlying domain-domain interaction. Toy scenarios that would result in the three possible rewiring events predicted by Splitpea are illustrated in B.

In the event that there are multiple exons within the same protein domain, we attribute the *minimum* ∆*ψ* value to the entire protein domain. The underlying assumption here is that loss of any portion of a particular protein domain may potentially negatively impact the protein domain’s downstream capacity to interact with other domains. Splitpea then determines the directionality of change based on whether or not there is consistency across the changing domains. In the event that there are mixed exon changes, the directionality is labeled as “chaos,” or undetermined (Fig. 1B). The weight of the edge is calculated as the mean domain-level ∆*ψ* values. Essentially, the following pseudocode describes the crux of Splitpea’s algorithm for a given sample with a set of exons with associated ∆*ψ* values:

**for** each PPI between *g*_*u*_, *g*_*v*_ **do**

Ψ^(*u*)^ := significant exons for gene u

Ψ^(*v*)^ := significant exons for gene v

*D*^(*u*)^ := *{d*_*u*_|*d*_*u*_ ∈ *g*_*u*_, ∃ exon_*i*_ s.t. exon_*i*_ ∈ Ψ^(*u*)^ & exon_*i*_ ∈ *d*_*u*_*}*

*D*^(*v*)^ := *{d*_*v*_|*d*_*v*_ ∈ *g*_*v*_, ∃ exon_*i*_ s.t. exon_*i*_ ∈ Ψ^(*v*)^ & exon_*i*_ ∈ *d*_*v*_*}*

*w*_*uv*_ := network rewiring edge weight between *g*_*u*_, *g*_*v*_

*δ*_*uv*_ := direction classification of network rewiring between *g*_*u*_, *g*_*v*_

**for** each DDI between *d*_*u*_ ∈ *D*^(*u*)^, *d*_*v*_ ∈ *D*^(*v*)^ **do**

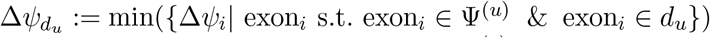

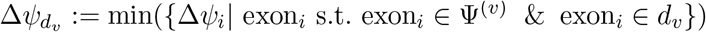

**if**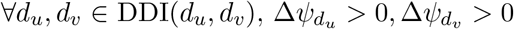 **then**

*δ*_*uv*_ = positive

**else if** ∀*d*_*u*_, *d*_*v*_ ∈ DDI(*d*_*u*_, *d*_*v*_), 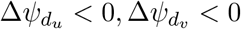, **then**

*δ*_*uv*_ = negative

**else**

*δ*_*uv*_ = chaos

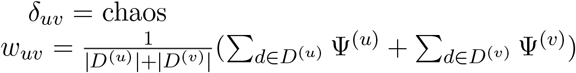

**return** *w*_*uv*_, *δ*_*uv*_

*w*_*uv*_ and *δ*_*uv*_ are reported as long as Ψ^(*u*)^ *or* Ψ^(*v*)^ is non-empty. Please note that the *w*_*uv*_ calculation only includes domains that have a DDI that is considered to be mediating the PPI between *g*_*u*_, *g*_*v*_. For readability, the equation above omits the removal of non-DDI pairs.

### 2.5 Consensus network

The main factor to consider when aggregating several sample-specific Splitpea networks into a consensus network is whether the directionality of edges agree. Thus, a “positive” consensus network and “negative” consensus network are built separately. “Chaos” edges are ignored since they are of ambiguous state. For each consensus network, two factors are considered for the edge weight: the sum of the original edge weights *w*_*uv*_ and how many networks support the same directionality *δ*_*uv*_. The downstream analysis with each consensus network focuses on the largest connected component. As is common in biological networks, we found that the largest connected component covers the majority of the edges of the complete consensus network (breast cancer: 96.4% edges retained in negative consensus, 89.5% edges retained in positive consensus; pancreatic cancer: 96.1% edges retained in negative consensus, 88.8% edges retained in positive consensus).

### 2.6 Network embedding and clustering

To enable network clustering and other downstream uses of the Splitpea patient-specific networks, we created whole graph level embeddings. Here, we chose to focus only on potential gain-of-interaction edges and first filtered each patient-specific network accordingly. Taking the largest connected component, we applied the FEATHER^41^ algorithm from the KarateClub NetworkX extension library^42^ to generate an embedding for each network.

We clustered the resulting embeddings for each cancer type using hierarchical density-based clustering (HDBSCAN)^43^ with minimum cluster sizes of 10. Clustering results were generally robust to the choice of the minimum cluster size parameter; 10 was chosen for downstream interpretability (and we would consider samples with fewer neighbors as outliers). Final plots were produced using principal component analysis (PCA), plotting all embeddings by their first two components.

## 3. Results

### 3.1 Quantifying splicing changes in pancreatic and breast tumors

In total, we collected data from TCGA covering 177 pancreatic primary tumors and 1,088 breast primary tumors, together with 192 normal pancreatic tissue and 218 normal breast tissue samples from GTEx that were used as a reference distribution of normal splicing variation for each respective cancer type. With these data, we calculated a ∆*ψ* value corresponding to the change in exon splicing in each tumor sample relative to its normal tissue background, resulting in ∆*ψ* estimates for a total of 139,661 unique exons across all breast cancer samples and 98,761 unique exons across the pancreatic cancer samples. Furthermore, we calculated an accompanying p-value that compares how extreme the observed *ψ* value for each exon in each cancer sample is relative to the corresponding background distribution of *ψ* values for normal tissue samples (see Methods).

### 3.2 ∆*ψ values primarily reflect primary diagnoses*

We then clustered the ∆*ψ* matrices for each tumor type and checked whether they corresponded to relevant clinical and pathological tumor features for both breast cancer (Fig. 2A) (pam50 subtypes, diagnosed type, pathologic stage, and age) and pancreatic cancer (Fig. 2B) (site of origin, diagnosed type, pathologic stage, age, and sex). While the majority of clinical features are not meaningfully clustered with ∆*ψ* values, we do observe that the most unique patient cluster for pancreatic cancer (far right columns in Fig. 2B) are all pancreatic neuroendocrine tumors. Neuroendocrine tumors are a rare subset of pancreatic cancers that originate not in the cells of the pancreas but in neuroendocrine cells. Interestingly, this cell type has commonality with neurons which are known to undergo more splicing changes.^44^ For breast cancer, we see some clustering of lobular carcinomas (red cluster in “type” bar Fig. 2A), but otherwise do not see obvious patterns of clinical or pathological separation with ∆*ψ* values alone.

**Fig. 2.**
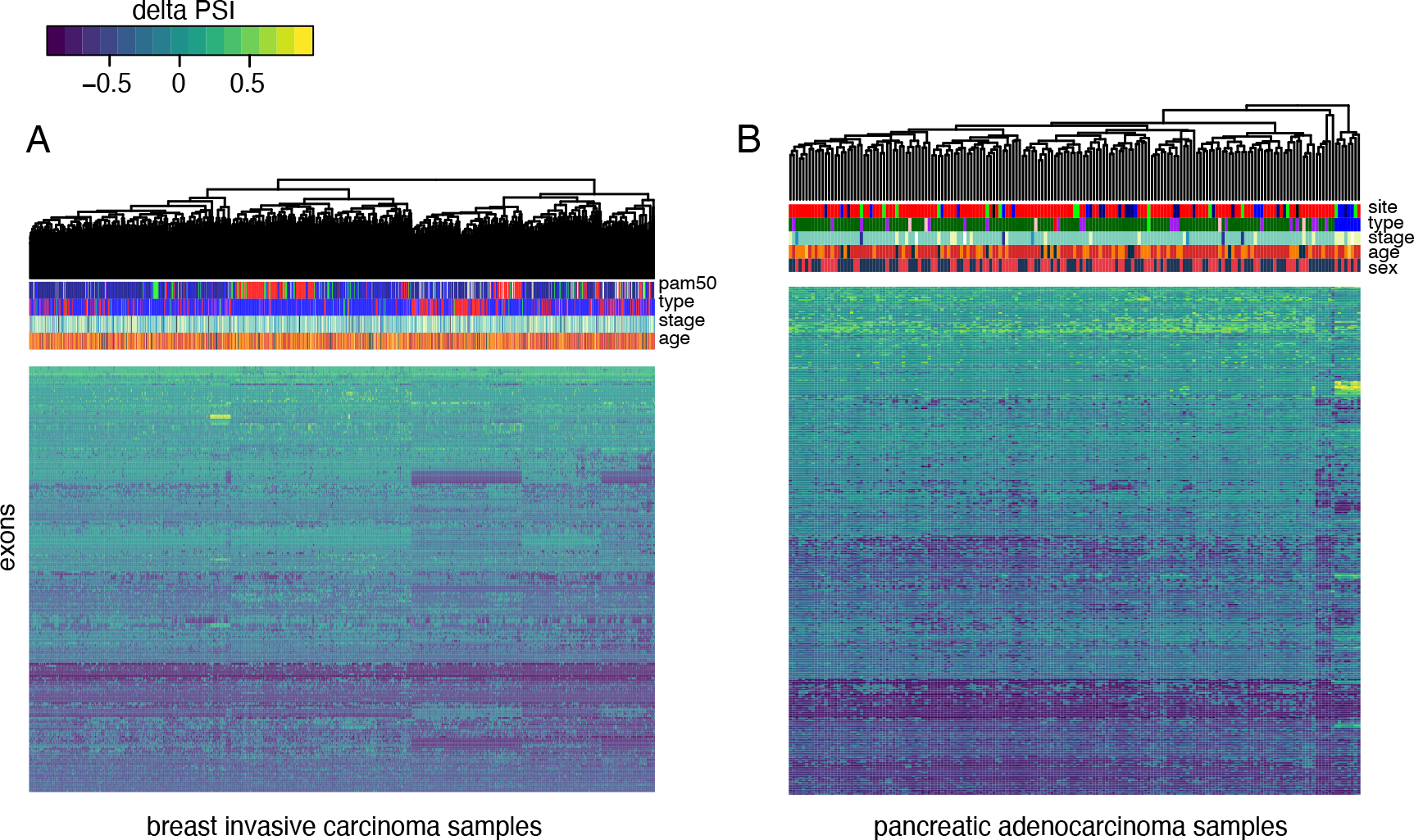
Clustering on ∆ψ values. We cluster the ∆*ψ* values showing different sample groups for different spliced exons. Heatmaps depict splicing changes relative to average normal tissue background. Bar columns show known clinical information about each sample. In general, there are more subgroup level exon changes for breast cancer, (A) but these are not strongly correlated with any clinical variable. In pancreatic cancer, a small subset of neuroendocrine samples (B, dark blue) share similar splicing patterns. All other samples do not have obvious meaningful structure.

### 3.3 Quantifying rewired protein-protein interactions for pancreatic and breast tumors

We applied Splitpea to build patient-specific rewired PPI networks for 177 pancreatic and 1,088 breast primary tumor samples. Each PPI network contains three types of edges (gain, loss, or chaotic (mixed)) based on how underlying splicing changes may affect the individual protein-protein interaction (Fig. 3). In general, most splicing changes cause potential loss of protein interactions, though breast cancer had relatively fewer loss of edges proportionally on average (76% edges) than pancreatic cancer (84% of edges). Chaos (mixed) edges, where domain interactions have inconsistent directions per protein are relatively uncommon and comprise on average less than 2% of total edges for pancreatic and breast cancer. Between the two cancer types, breast cancer has more potential gain-of-interaction edges and a lower proportion of potential lost edges relative to pancreatic cancer. Interestingly, there is also more variability across edge types per sample in breast cancer samples.

**Fig. 3.**
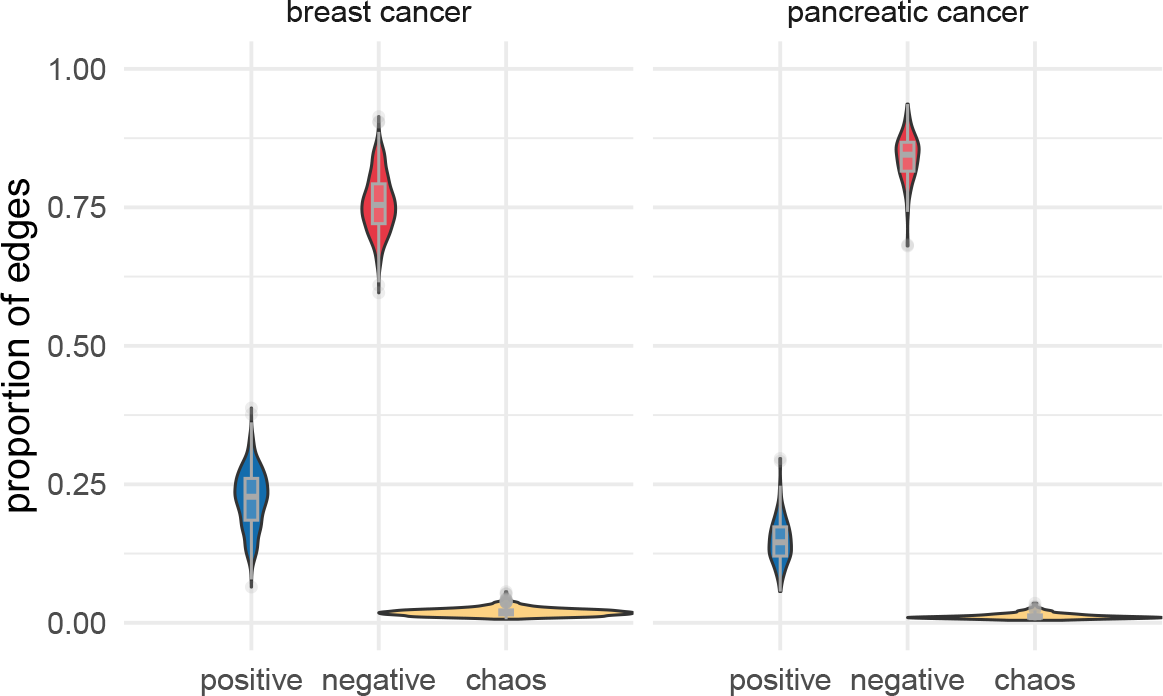
Proportion of relative gain and loss in edges across breast cancer and pancreatic cancer samples. Breast cancer samples have proportionally more “gain of interactions” than pancreatic cancer samples, but in both cancer types, interaction loss is much more prevalent. For each TCGA cancer sample, the proportion of edges gained versus lost is calculated using the total number of edges in the largest connected component of the entire Splitpea rewired network (both directions) as the denominator. To be conservative, the number of edges retained in the largest connected components for the gain-only subnetwork and loss-only subnetworks are used as numerators.

Looking at individual patient networks (Fig. 4), we can see potential hubs and protein clusters that undergo extensive remodeling. In Fig. 4A, we show an example of one pancreatic tumor network with the most remodeling changes in the oncogene, RAB35, proto-oncogenes, HRAS and FYN, the signaling protein, MAPK3, the cell cycle and growth genes, NEDD8 and PRKAA1, among others. Breast cancer patient-specific networks have a different topology (Fig. 4C), though there is also overlap of proto-oncogenes HRAS and FYN.

**Fig. 4.**
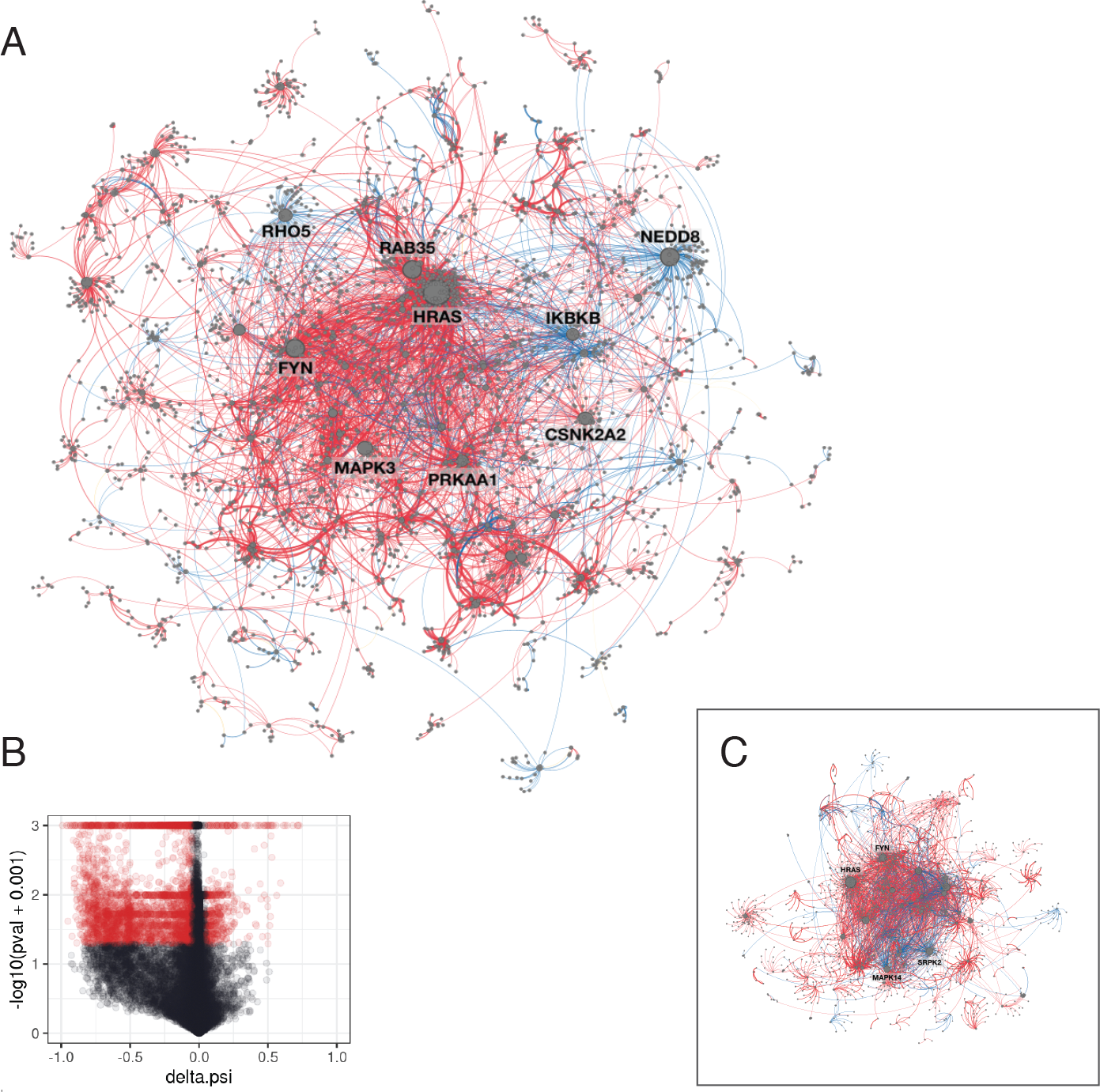
Patient specific rewired networks. Here, we show two sample network outputs from Splitpea and the accompanying exon value cutoff. The large network (A) depicts pancreatic patient sample (TCGA-HZ-7918-01A-11R-2156-07), with edge losses in red and gains in blue. The corresponding volcano plot is shown in (B), where exons with significant 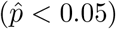 as well as absolute change (|∆*ψ*| *>* 0.05) are shown in red. Box (C) shows a patient-specific network for an example breast cancer sample, TCGA-BH-A0BG-01A-11R-A115-07, which exhibits a very different topology from the pancreatic sample in A.

### 3.4 A consensus network of changes across breast cancer patients

While patient-specific networks highlight network rewiring at the level of individual tumor samples, we also sought to look for more general cancer level patterns of PPI rewiring. Towards this end, we assembled a consensus rewiring network for breast cancer by taking splicing rewiring events conserved across 80% of patient samples and assembling a meta-network of these events. Edges were only preserved when their type (gain, loss) was consistent. Chaos edges were not included in the consensus network. Naturally, as the threshold increases, the number of genes preserved in the network decreases (Fig. 5A). Interestingly, up through the 80% threshold, gained edges are relatively more consistently preserved (Fig. 5B). Visualizing the breast cancer consensus network (Fig. 5C) revealed that the most gained interaction involved the gene, FKBP5, which is an immune regulator responsible for protein trafficking and folding. This protein has been studied in breast cancer for its various hormone receptor signaling functions.^45^

**Fig. 5.**
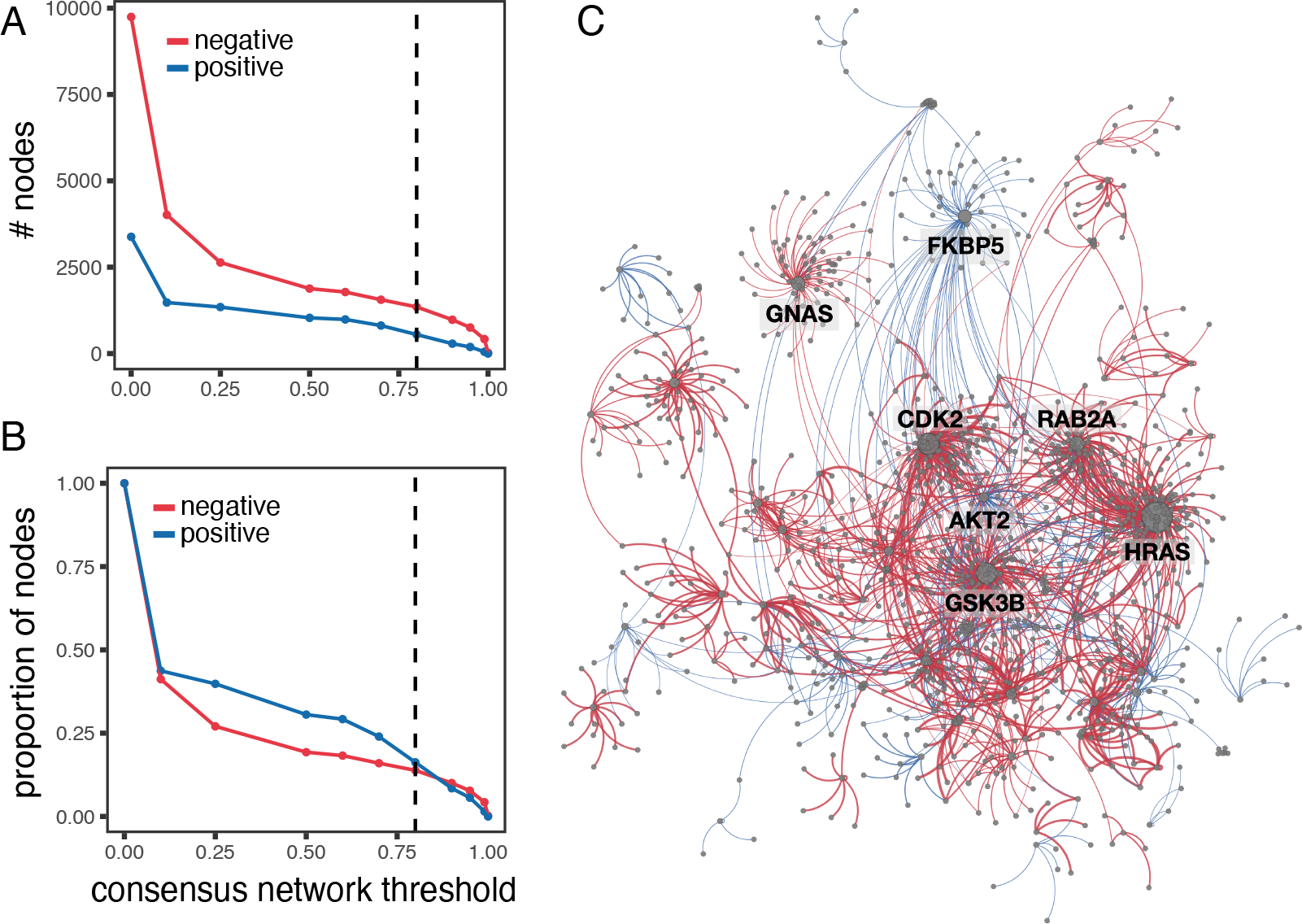
Meta-network of breast cancer patients. The line graphs show the number of nodes preserved for different consensus thresholds (A) or the proportion of nodes relative to the non-thresholded consensus network (B) for edge loss (negative, red) and edge gain (positive, blue) events. The dashed line in both graphs denotes a threshold of 80%, corresponding to the visualization of the consensus network of splicing rewiring events conserved across 80% of breast cancer patient samples (C, red: edge loss; blue: edge gain).

### 3.5 Network clusters reveal novel patient subgroups

The patient-specific networks generated by Splitpea have many downstream applications, especially when the networks are used as features for other machine learning tasks. Here, we demonstrate their utility by finding patient subgroups across both breast and pancreatic cancer when the networks are clustered (Fig. 6). Specifically, we use a state-of-the-art graph embedding method, FEATHER,^41^ which calculates characteristic functions using different random walk weights for node features, but any graph embedding method could be used for this type of analysis. For each cancer type, we clustered the network embeddings using HDBSCAN (see Methods). Interestingly, three distinct groups emerged across the cancer types (Fig. 6A). The dominant source of variation across the networks is the gain or loss of PPIs involving KRAS (Fig. 6B). Mutations in KRAS are known to affect subgroups of both pancreatic and breast cancer^46^ with ties to prognosis. It is possible that splicing changes in interacting partner genes also induce changes to KRAS that may have yet unknown interaction effects with these somatic mutations, highlighting the potential of Splitpea to find additional disease subtypes. Furthermore, other interesting cancer drivers have distinct patterns of gains and losses, including RAB5A, which appears to have PPI gains in the BRCA outliers, and IKBKB, which is enriched for gains in the predominantly pancreatic cancer cluster 3.

**Fig. 6.**
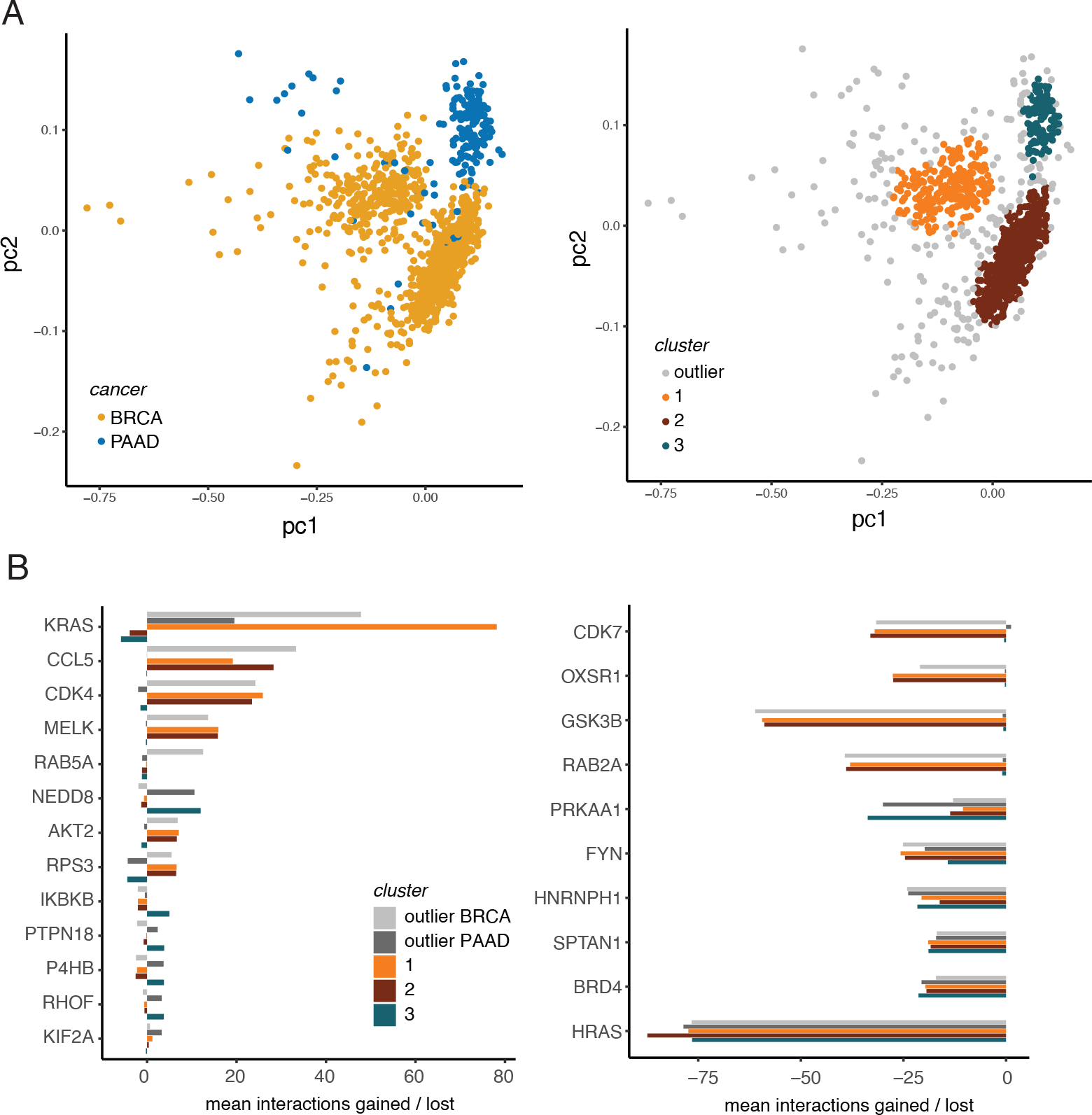
Splitpea networks cluster into distinct subgroups. (A) PCA plots of graph embeddings of each patient-specific Splitpea network, with samples colored by either cancer type (left) or cluster (right). Clusters were assigned using HDBSCAN, with outliers colored in grey. (B) For each cluster, the top nodes undergoing the most changes (mean interactions gained or lost) were also identified. The bar graphs are roughly separated by genes that have the most gain of interactions (left) versus those that have primarily losses (right). Interestingly, the main variation captured in PC1 seems to be defined by networks that change in KRAS. Other cancer driver genes also undergo distinct patterns of gains and losses that drive clustering patterns.

## 4. Discussion and conclusion

We present a new method, Splitpea, for characterizing protein-protein network rewiring events. Splitpea is flexible and can be applied with different background contexts to highlight splicing changes between a disease and relevant background context of interest. We applied Splitpea to breast and pancreatic cancer samples to highlight the potential of Splitpea to find new and relevant cancer biology, both on an individual patient sample level and more broadly across samples of a single tumor type. To our knowledge, Splitpea is the first systematic method for identifying both potential gains in addition to PPIs lost for individual experimental samples.

Splitpea makes heavy use of existing knowledge of protein-protein interactions. Because of this, our method is inherently limited by the availability of known PPIs (which are largely incomplete), as well as DDIs, which are even less complete. As more of these are experimentally characterized, Splitpea will continue to improve, capturing more accurate and comprehensive sets of network rewiring events. Since we wrote Splitpea to be modular, updates to PPIs and DDIs can be easily integrated once they become available. Specifically, study bias is a well-reported issue in PPIs, and thus there is a large amount of overlap between well-studied nodes (including many cancer driver genes) with nodes of high degree in PPI networks, and given the dependency of Splitpea on reported PPIs, this also affects our results. As more systematic experimental PPI screens and more reliable PPI predictions become available, we can also readily adapt Splitpea.

We have only scratched the surface of cancer biology here. In our initial exploration of breast and pancreatic cancer, we have discovered subgroups and outliers within each cancer type that can be characterized by different network hubs. We believe this merits more thorough exploration, as it may carry important implications for precision medicine efforts. Beyond this, it will also be interesting to apply Splitpea to more cancer types and look for pan-cancer conservation patterns.

## Acknowledgments

This work was supported by the Cancer Prevention & Research Institute of Texas (RR190065). VY is a CPRIT Scholar in Cancer Research.

## Notes

https://github.com/ylaboratory/splitpea

### Competing Interest Statement

The authors have declared no competing interest.

### Summary of Updates

updates for camera ready version

